## Supplemental Information for "Extracellular prostaglandins E1 and E2 and inflammatory cytokines are regulated by the senescence program in potentially premalignant oral keratinocytes"

**Supplementary Methods**

**Cell Lines used in the study**

The characteristics, culture methods and properties of the LR- and HRPPOL line used in the study have been published recently and in previous publications [1-4]. Oral keratinocytes from the floor of the mouth were immortalized by the group of James Rheinwald. Oral keratinocytes floor of mouth line 6 (OKF6) was immortalized by the ectopic expression of the telomerase catalytic subunit *TERT (*OKF6/TERT-1*)* hitherto referred to as OKF6 [5]. Oral keratinocytes floor of mouth line 64 (OKF4/CDK4R/P53DD/TERT), hitherto referred to as OKF4, was immortalized by the defined genetic elements Cdk4 mutant, the dominant-negative p53DD peptide and ectopic expression of *TERT* OKF4/CDK4R/P53DD/TERT [6].

**Conditioned medium collection**

To achieve confluence in 72 hours the cells were disaggregated, the trypsin/EDTA neutralized, counted on a haemocytometer pelleted by centrifugation at 300g and the amount of protein in each cell pellet was measured and the results expressed in pg/ml medium/mg cell protein. The cell densities plated to achieve confluence in 72 hours are as follow s: OKF4, 1.1. x 10^5^/square cm ; OKF6, 1 3.28 x 10^5^; D19, 1.44 x 10^5^; D35, 1.6 x 10^5^; DOK, 7.2 x 10^4^ NHOK810, 4 x 10^4^; D6, 5.5 x 10^4^; D9, 1.06 x 10^5^; E4, 3.2 x 10^4^; D25, 8 x 10^4^; D4, 5.5 x 10^3^; D20, 8.3 x 10^5^; D34, 6.67 x 10^4^ and D17, 1.33 x 10^5^ .

**Knockdown of p53, COX-1 and COX-2 in OKF6**

The target sequences of the siRNAs are detailed in Supplementary Table S1.

**Supplementary Table S1 si RNA target sequences**

| **siRNA** | **Target sequences** |
| --- | --- |
| ON-TARGETplus SMARTpool Human TP53 siRNA | 5’-GAAATTTGCGTGTGGAGTA-3’  5’-GTGCAGCTGTGGGTTGATT-3’  5’-GCAGTCAGATCCTAGCGTC-3’  5’-GGAGAATATTTCACCCTTC-3’ |
| ON-TARGETplus SMARTpool Human PTGS1 siRNA | 5’-CCACAUUUAUGGAGACAAU-3’  5’-GAAUCCCUGUUGUUACUAU-3’  5’-CGAGGUGGGCUUUAACAUU-3’  5’-UCAAGGGUCUCCUAGGGAA-3’ |
| ON-TARGETplus SMARTpool Human PTGS2 siRNA | 5’-GGACUUAUGGGUAAUGUUA-3’  5’-GAUAAUUGAUGGAGAGAUG-3’  5’-GUGAAACUCUGGCUAGACA-3’  5’-CGAAAUGCAAUUAUGAGUU-3’ |
| ON-TARGETplus SMARTpool Human CDKN2A siRNA | 5’-GATCATCAGTCACCGAAGG-3’  5’-AAACACCGCTTCTGCCTTT-3’  5’-TAACGTAGATATATGCCTT-3’  5’-CAGAACCAAAGCTCAAATA-3’ |
| ON-TARGETplus Non-targeting pool | 5’-TGGTTTACATGTCGACTAA-3’  5’-TGGTTTACATGTTGTGTGA-3’  5’-TGGTTTACATGTTTTCTGA-3’  5’-TGGTTTACATGTTTTCCTA-3’ |

Several rounds of optimisation were done to determine the optimised cell density, final TP53 siRNA concentrations, volume of lipid complex (DharmaFECT transfection reagent) and transfection incubation time. TP53 was optimised with cell density 2.5x10^5^ in each well of a six well plate, 100nM final siRNA concentration, and 5µl of DharmaFECT transfection reagent and 24-hours transfection incubation time. The cells were plated in antibiotic-free complete medium (DMEM with 10% FetalClone® II foetal bovine serum, 2mM L-glutamine and 0.4µg/ml hydrocortisone). The cells were incubated at 37^o^C overnight. The transfection medium was replaced with complete medium containing antibiotics after 24 hours to reduce cytotoxicity. The cells were incubated for an additional 24 hours before being harvested for western blot analysis and quantitation of the level of knock down as well as conditioned medium collection.

**Western blotting**

**Supplementary Table S2 Details of the antibodies and optimised dilutions.**

| **Antibody** | **Clonality** | **Species** | **Optimised dilution** | **Sources** |
| --- | --- | --- | --- | --- |
| Anti-p16INK4a (ab81278) | Monoclonal | Rabbit | 1:2000 | Abcam, Cambridge, UK |
| P16-INK4A (10883-1-AP) | Polyclonal | Rabbit | 1:5000 | Proteintech, Manchester, UK |
| Anti-p53 (ab1101) | Monoclonal | Mouse | 1:1000 | Abcam, Cambridge, UK |
| Phospho-p53 (Ser15) (#9286) | Monoclonal | Mouse | 1:500 | Cell Signaling Technology, Beverly, USA |
| Anti-MCM7 [EP1974Y] (ab52489) | Monoclonal | Rabbit | 1:10000 | Abcam, Cambridge, UK |
| Anti-SIRT1 [E104] (ab32441) | Monoclonal | Rabbit | 1:20000 | Abcam, Cambridge, UK |
| Anti-Chk2 [EPR4325] (ab109413) | Monoclonal | Rabbit | 1:50000 | Abcam, Cambridge, UK |
| Anti-Chk2 (phosphor T68) [Y171] (ab32148) | Monoclonal | Rabbit | 1:1000 | Abcam, Cambridge, UK |
| Anti-ATM [Y170] (ab32420) | Monoclonal | Rabbit | 1:10000 | Abcam, Cambridge, UK |
| Anti-ATM (phosphor S1981) [EP1890Y] (ab81292) | Monoclonal | Rabbit | 1:50000 | Abcam, Cambridge, UK |
| EP1 Receptor (101740) | Polyclonal | Rabbit | 1:200 | Cayman Chemical, Michigan, USA |
| Anti-Prostaglandin E Receptor EP1 (ab183073) | Polyclonal | Rabbit | 1:450 | Abcam, Cambridge, UK |
| EP2 Receptor (101750) | Polyclonal | Rabbit | 1:500 | Cayman Chemical, Michigan, USA |
| Anti-Prostaglandin E Receptor EP2 [EPR8030(B)] (ab167171) | Monoclonal | Rabbit | 1:1000 | Abcam, Cambridge, UK |
| Anti-PTGER3 (ab117998) | Polyclonal | Rabbit | 1:2000 | Abcam, Cambridge, UK |
| Anti-Prostaglandin E Receptor EP4 (ab45295) | Polyclonal | Rabbit | 1:1000 | Abcam, Cambridge, UK |
| Anti-COX1 / Cyclooxygenase 1 [EPR5866] (ab109025) | Monoclonal | Rabbit | 1:1000 | Abcam, Cambridge, UK |
| Anti-COX2 / Cyclooxygenase 2 [EPR8588] (ab151571) | Monoclonal | Rabbit | 1:1000 | Abcam, Cambridge, UK |
| Anti-Notch 1 (ab87982) | Monoclonal | Mouse | 1:500 | Abcam, Cambridge, UK |
| Cleaved Notch1 (Val1744) (D3B8) (#4147) | Monoclonal | Rabbit | 1:1000 | Cell Signaling Technology, Beverly, USA |
| Anti-β-actin (ab8227) | Polyclonal | Rabbit | 1:20000 | Abcam, Cambridge, UK |
| GAPDH (14C10) (#2118) | Monoclonal | Rabbit | 1:4000 | Cell Signaling Technology, Beverly, USA |
| Anti-CREB (phosphor S133) [E113] (ab32096) | Monoclonal | Rabbit | 1:5000 | Abcam, Cambridge, UK |
| CD44 (156-3C11) (#3570) | Monoclonal | Mouse | 1:1000 | Cell Signaling Technology, Beverly, USA |
| Anti-Integrin alpha 6 [EPR18124] (ab181551) | Monoclonal | Rabbit | 1:2000 | Abcam, Cambridge, UK |
| Anti-FADS1 [EPR6898] (ab126706) | Monoclonal | Rabbit | 1:1000 | Abcam, Cambridge, UK |

**Supplementary Table S2:** The dilution used for each antibody was optimised to obtain clear band on the positive controls. Wherever possible monoclonal antibodies were used to reduce unspecific binding of the protein of interest.

**Supplementary Table S3 Details of the antibodies and the positive and negative controls.**

| **Antibody** | **Positive Control** | **Negative Control** |
| --- | --- | --- |
| Anti-p16INK4a (ab81278) | HeLa | BICR3p16^INK4A^-/-  P14^ARF^/p15^INK4B^ + |
| P16-INK4A (10883-1-AP) | HeLa | BICR3p16^INK4A^-/-  P14^ARF^/p15^INK4B^ + |
| Anti-p53 (ab1101) | BICR3 Mis-sense mutation codon 282 /HEK293 | BICR6 Stop codon 192 |
| Phospho-p53 (Ser15) (#9286) | Fibroblasts irradiated with 10Gy Gamma rays and cultured for8 hours | BICR6 Stop codon 192 |
| Anti-MCM7 [EP1974Y] (ab52489) | Senescent fibroblasts | Growing fibroblasts |
| **Antibody** | **Positive Control** | **Negative Control** |
| Anti-SIRT1 [E104] (ab32441) | Senescent Fibroblasts | Growing/quiescent fibroblasts |
| Anti-Chk2 [EPR4325] (ab109413) | Fibroblasts irradiated with 10Gy Gamma rays and cultured for8 hours |  |
| Anti-Chk2 (phosphor T68) [Y171] (ab32148) | Fibroblasts irradiated with 10Gy Gamma rays and cultured for8 hours |  |
| Anti-ATM [Y170] (ab32420) | Fibroblasts irradiated with 10Gy Gamma rays and cultured for8 hours | ATM-deficient human keratinocytes from patient AT5BI |
| Anti-ATM (phosphor S1981) [EP1890Y] (ab81292) | Fibroblasts irradiated with 10Gy Gamma rays and cultured for8 hours | ATM-deficient human keratinocytes from patient AT5BI |
| EP1 Receptor (101740) | Previously validated | Previously validated |
| Anti-Prostaglandin E Receptor EP1 (ab183073) | Previously validated | Previously validated |
| EP2 Receptor (101750) | Previously validated | Previously validated |
| Anti-Prostaglandin E Receptor EP2 [EPR8030(B)] (ab167171) | Previously validated | Previously validated |
| Anti-PTGER3 (ab117998) | Previously validated | Previously validated |
| Anti-Prostaglandin E Receptor EP4 (ab45295) | Previously validated | Previously validated |
| Anti-COX1 / Cyclooxygenase 1 [EPR5866] (ab109025) | Mock/Scrambled siRNA transfected keratinocytes | Smartpool COX1 siRNA transfected keratinocytes |
| Anti-COX2 / Cyclooxygenase 2 [EPR8588] (ab151571) | Mock/Scrambled siRNA transfected keratinocytes | Smartpool COX2 siRNA transfected keratinocytes |
| Anti-Notch 1 (ab87982) | Previously validated | Previously validated |
| Cleaved Notch1 (Val1744) (D3B8) (#4147) | Previously validated | Previously validated |
| Anti-β-actin (ab8227) | Previously validated | Previously validated |
| GAPDH (14C10) (#2118) | Previously validated | Previously validated |
| Anti-CREB (phosphor S133) [E113] (ab32096) | Previously validated | Previously validated |
| CD44 (156-3C11) (#3570) | Previously validated | Previously validated |
| Anti-Integrin alpha 6 [EPR18124] (ab181551) | Previously validated | Previously validated |

**Western blotting reagents**

RIPA buffer (Sigma, Dorset, UK)

cOmplete™ Mini EDTA-free Protease Inhibitor Cocktail (Roche, Burgess Hill, UK

Phosphatase inhibitor (PhosSTOP; Roche, Mannheim, Germany)

NuPAGE^TM^ NOVEX^TM^ SDS sodium dodecyl sulphate polyacrylamide gels. (Bis-Tris pre-cast, ThermoFisher Scientific, Dartford, UK).

Skimmed milk (Marvel, Ashford, UK).

Tris Buffer Saline and Tween 20 (TBS-T) (Sigma-Aldrich, Dorset, UK).

PIERCE^TM^ ECL Western Blotting Substrate (ThermoFisher Scientific, Dartford, UK).

Amersham ECL Prime Western Blotting Detection Reagent (GE Healthcare, Buckinghamshire, UK).

SuperSignal® West Femto Maximum Sensitivity Substrate (Thermo Scientific, Rockford, USA).

Amersham Hyperfilm ECL (GE Healthcare, Buckinghamshire, UK).

Film developer machine (Konica Minolta, Banbury, UK).

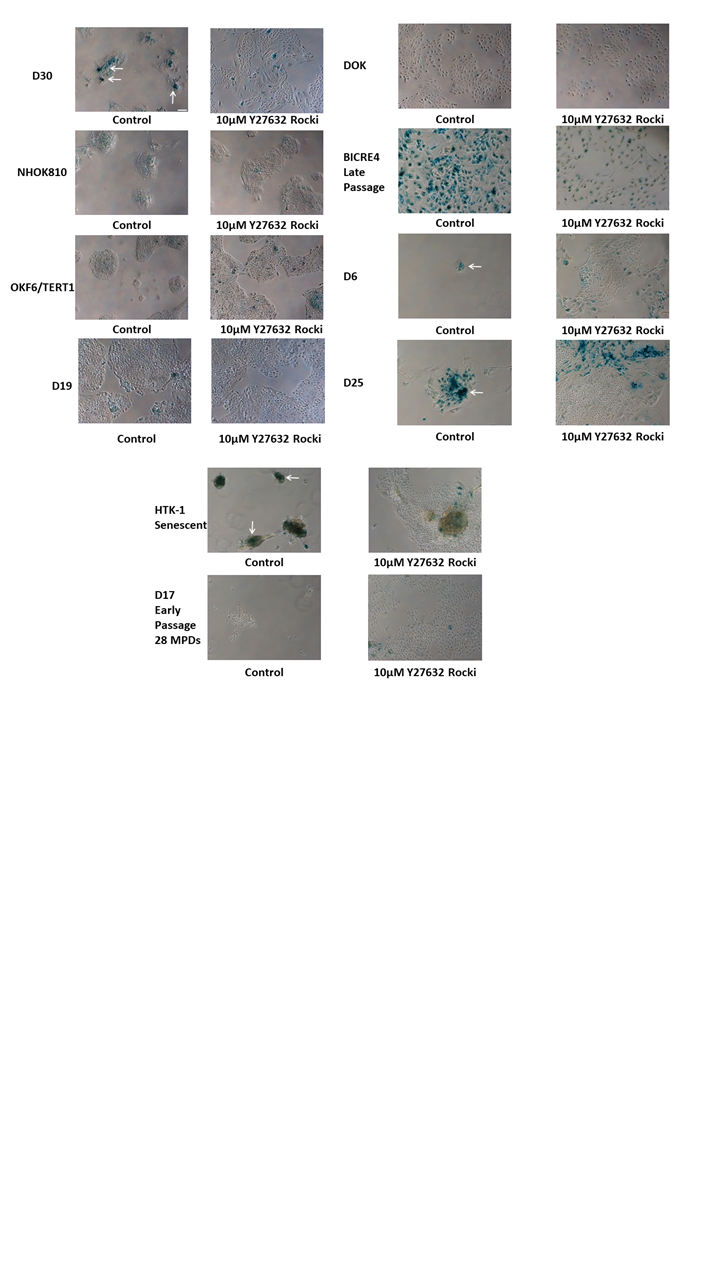
**Supplementary Figure S1: SA-βGal staining of MPPOL and IPPOL keratinocytes and the effect of ROCKi.**

The Figure shows the SA-βGal staining in control normal, MPPOL,

IPPOL and normal immortal OKF6 keratinocytes. Note the location of the blue cells indicated in the control and MPPOL cultures indicated by the white arrows in the centers of the colonies. These positively stained areas largely disappear following 2 weeks culture in ROCKi and the cultures continue to proliferate. The blue cells in the D25 ROCKi panel are lethally irradiated (senescent) 3T3 fibroblasts that have not been removed by the EDTA treatment. Bar = 100µm see D30 control panel).
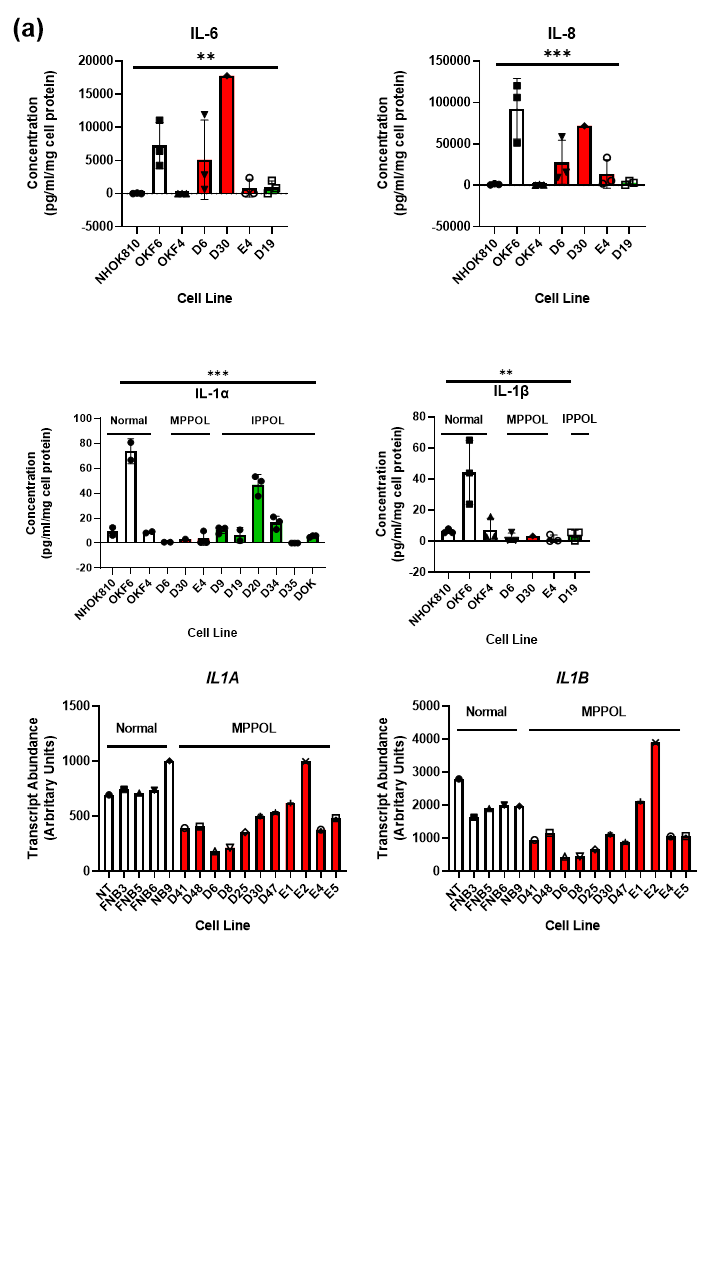

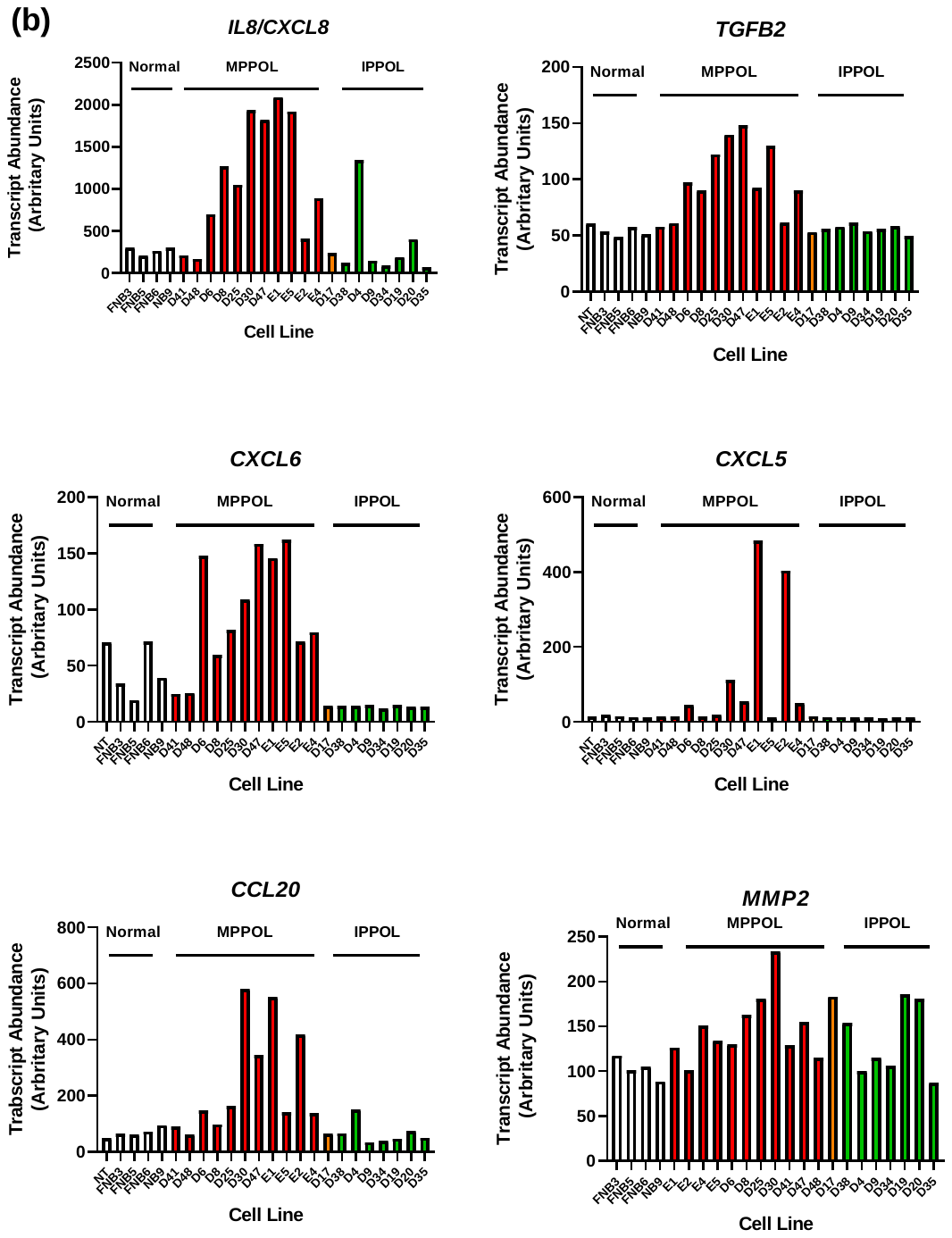

**Supplementary Figure S2 Levels of secreted SASP cytokine protein and SASP transcript in normal, MPPOL and IPPOL keratinocytes.**

(a) The top four panels show cytokine protein levels in normal NHOK810 and normal immortal OKF4 and OKF6 keratinocytes (white bars) MPPOL keratinocytes D6, D30 and E4 (red bars) and IPPOL keratinocytes (green bars). The results are significantly different by one way ANOVA ** < 0.01 *** < 0.001 (n =3) and show an increase in extracellular IL-6 and IL-8 in all 3 MPPOL lines over NHOK810 but if anything the same lines showed lower levels of IL-1α and IL-1β than normal. The bottom two panels show the levels of SASP transcript in a large panel of normal (n =5), MPPOL (n =11), D17 (MPPOL; p16^INK4A^-/-) and IPPOL (n =7) keratinocytes mined from reference [1]. Symbols the same as the top two panels except D17 bars are orange.
(b) Transcript abundance of SASP cytokines upregulated in MPPOL keratinocytes relative to normal. Data mined from reference [1]. Symbols are the same as in (a). The data show elevated SASP transcripts in MPPOL are generally revered in IPPOL keratinocytes.

**Supplementary Figure S3 ePGE regulation by DNA damage in keratinocytes.**

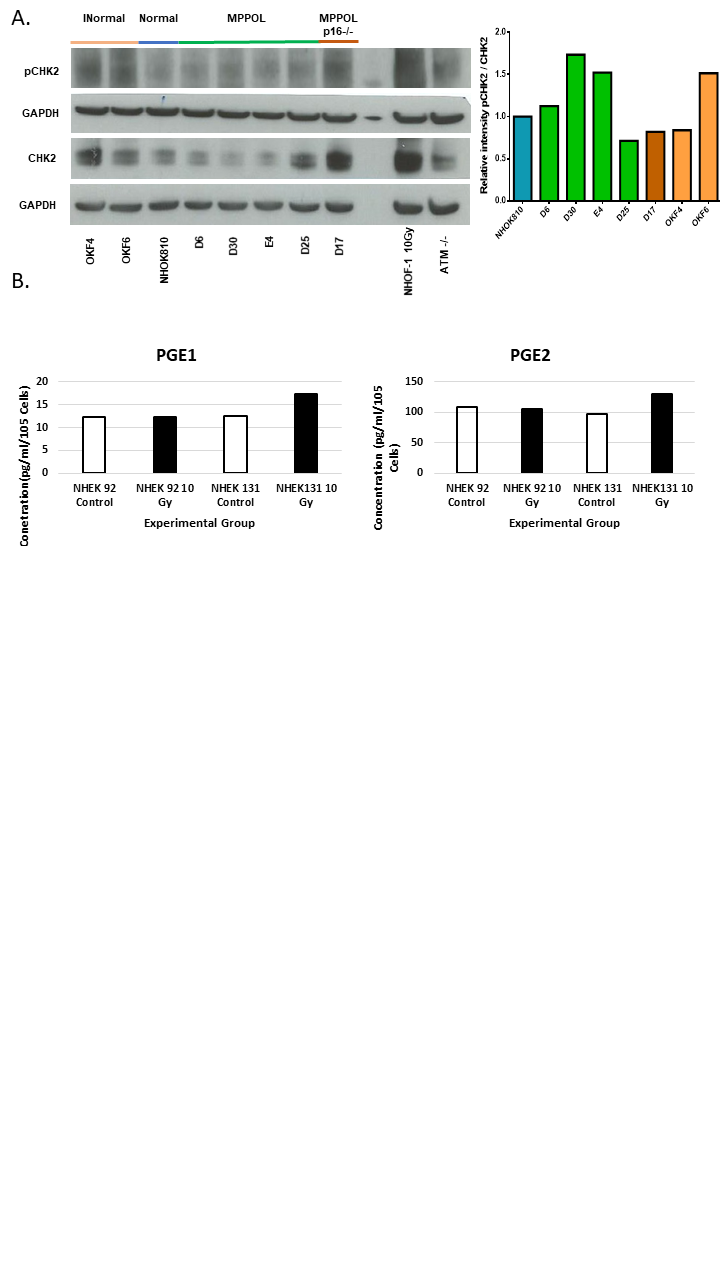

A. **CHK2 is not regulated in PPOL keratinocytes and is independent of ATM.** Western blot of (A) pCHK2 (phosphorylated at tyrosine residue T68) and total CHK2 in NHOK810, subset of the LRPPOL, OKF4, OKF6 and D17. Normal oral fibroblast irradiated with 10Gy of gamma rays (NHOF-1 10Gy) and cultured for 8 hours was used as a positive control for DDR protein phosphorylation (pCHK2 and CHK2) while normal keratinocytes from an ataxia telangiectasia patient (ATM-/-) were used as a control for ATM function. GAPDH was used as a loading control in all cases. All lanes contain equal amounts of protein (20µg), pCHK2 with histograms represent mean ratios of the protein levels relative to the total CHK2 and GAPDH loading control respectively. The figure represents data from single blot from protein extracts derived from one of the experiments in Figure 1.
B. **Measurements of prostaglandins (PGE1 and PGE2) in two commercial lines of epidermal keratinocytes (NHEK92 and NHEK131).** Both lines were irradiated at 10Gy of Gamma Rays to introduce irreparable DNA double strand breaks (IrrDSBs) and then cultured for 10 days to induce senescence. The white bars represent pre-irradiated lines while the black bars represent irradiated lines (10Gy). Concentration of the PGEs were normalised to the total cell count (x10^5^)/ml. All data is derived from a single experiment on two lines (n=1).

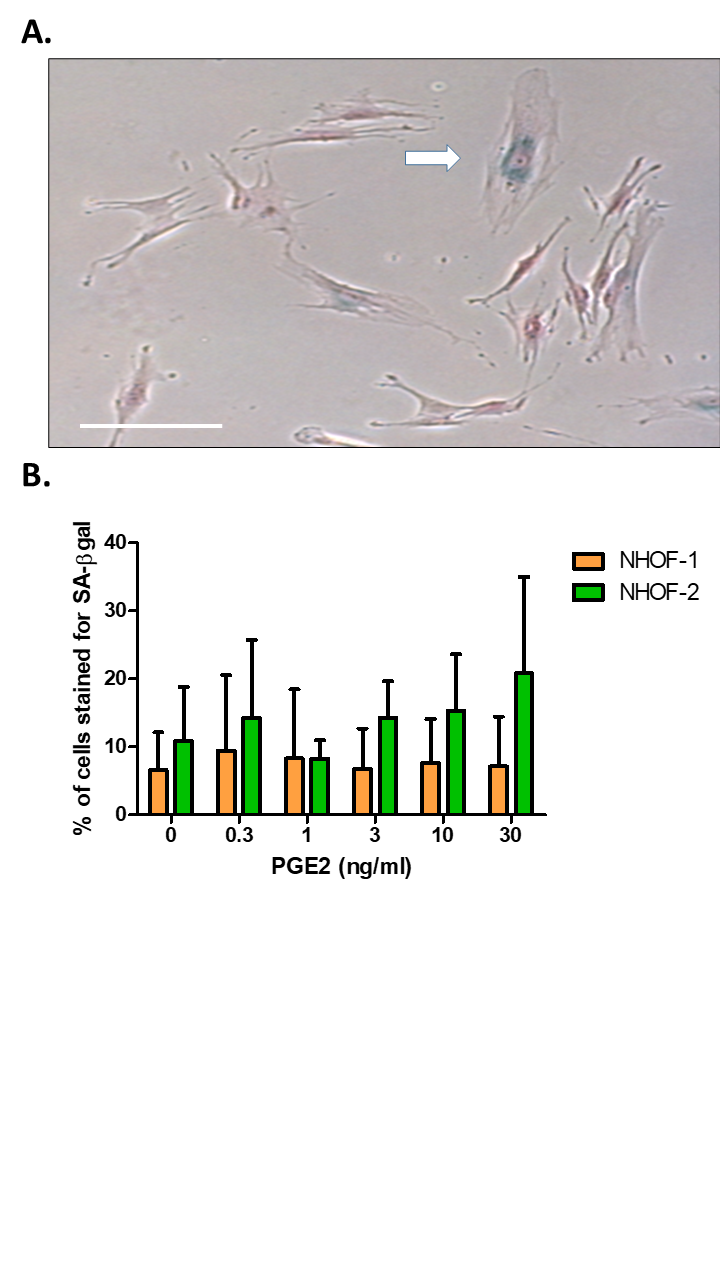
**Supplementary Figure S4: A.** Representative image of the senescence phenotype in oral fibroblast cells stained positive blue (as shown by the arrow) for SA-β-Gal activity upon treatment with PGE2. The bar in the figure represents 100 μm. **B.** Percentage of fibroblasts positive for SA-β-Gal activity – positive blue cells. The oral fibroblasts (NHOF-1 and NHOF-2) were exposed to different concentrations (0.3, 1, 3, 10 and 30 ng/ml) of PGE2 and stained for SA-β-Gal after a 24-hour exposure. The NHOF-1 and NHOF-2 were not treated with PGE2 in the control (0 ng/ml). SA-β-Gal positive blue cells were counted in 3 different microscopic fields for each of the PGE2 concentration exposure. The histograms represent the mean ± SD of the 9 counts (3 counts x 3 repeats) from the representative results of 3 independent experiments (n=3). T-tests were done to compare means of positive stained for SA-β-Gal from each concentration in both NHOF-1 and NHOF-2 against the control (0 ng/ml). No significant difference was found between the treated and untreated groups.

1.	Hunter, K. D.; Thurlow, J. K.; Fleming, J.; Drake, P. J.; Vass, J. K.; Kalna, G.; Higham, D. J.; Herzyk, P.; Macdonald, D. G.; Parkinson, E. K.; Harrison, P. R., Divergent routes to oral cancer. *Cancer Res* **2006,** *66*, 7405-13.

2.	Karen-Ng, L. P.; James, E. L.; Stephen, A.; Bennett, M. H.; Mycielska, M. E.; Parkinson, E. K., The Extracellular Metabolome Stratifies Low and High Risk Potentially Premalignant Oral Keratinocytes and Identifies Citrate as a Potential Non-Invasive Marker of Tumor Progression. *Cancers (Basel)* **2021,** *13*.

3.	McGregor, F.; Muntoni, A.; Fleming, J.; Brown, J.; Felix, D. H.; MacDonald, D. G.; Parkinson, E. K.; Harrison, P. R., Molecular changes associated with oral dysplasia progression and acquisition of immortality: potential for its reversal by 5-azacytidine. *Cancer Res* **2002,** *62*, 4757-66.

4.	Muntoni, A.; Fleming, J.; Gordon, K. E.; Hunter, K.; McGregor, F.; Parkinson, E. K.; Harrison, P. R., Senescing oral dysplasias are not immortalized by ectopic expression of hTERT alone without other molecular changes, such as loss of INK4A and/or retinoic acid receptor-beta: but p53 mutations are not necessarily required. *Oncogene* **2003,** *22*, 7804-8.

5.	Dickson, M. A.; Hahn, W. C.; Ino, Y.; Ronfard, V.; Wu, J. Y.; Weinberg, R. A.; Louis, D. N.; Li, F. P.; Rheinwald, J. G., Human keratinocytes that express hTERT and also bypass a p16(INK4a)-enforced mechanism that limits life span become immortal yet retain normal growth and differentiation characteristics. *Mol Cell Biol* **2000,** *20*, 1436-47.

6.	Rheinwald, J. G.; Hahn, W. C.; Ramsey, M. R.; Wu, J. Y.; Guo, Z.; Tsao, H.; De Luca, M.; Catricala, C.; O'Toole, K. M., A two-stage, p16(INK4A)- and p53-dependent keratinocyte senescence mechanism that limits replicative potential independent of telomere status. *Mol Cell Biol* **2002,** *22*, 5157-72.
